# The hidden life of *Xylella*: Mining the NCBI Sequence Read Archive reveals potential new species, host plants and infected areas for this elusive bacterial plant pathogen

**DOI:** 10.1101/2025.04.18.649540

**Authors:** Martial Briand, Marie-Agnès Jacques, Jessica Dittmer

**Author notes:** Address correspondence to: Martial Briand.

## Abstract

New crop disease outbreaks can occur when phytopathogenic bacterial pathogens acquire new traits, switch to a new host plant or are introduced into new geographic areas. Therefore, the rapid detection of a pathogen in a new environment and/or in a new host plant is essential to mitigate disease outbreaks. However, bacteria with a wide plant host range, many asymptomatic hosts and slow symptom development can sometimes remain unnoticed for years. This is the case for the vector-borne xylem-inhabiting bacteria *Xylella fastidiosa* known to infect hundreds of plant species worldwide and its sister species *X. taiwanensis*, currently thought to be restricted to Taiwan. To investigate whether the two *Xylella* species are already present in other parts of the world, potentially in unrecognised host species, we performed an in-depth data mining of raw sequence data available in the NCBI Sequence Read Archive. This led to the identification of 62 datasets from diverse plant and insect samples from around the world. Furthermore, nine draft and one circular *Xylella* genome could be assembled from these datasets. Our results reveal several potential new host plants and previously unrecognized infected areas in the Americas, Africa and southeast Asia. Moreover, the newly-assembled genomes represent several new strains of both *X. fastidiosa* and *X. taiwanensis* as well as an additional *Xylella* species infecting wild rice. Taken together, our work extends our knowledge on the genetic diversity, host range and global distribution of the genus *Xylella* and can orient surveillance programs towards new regions and host plants.

## INTRODUCTION

Crop diseases caused by phytopathogenic bacteria cause important economic losses worldwide. New disease outbreaks can occur when a pathogen acquires new traits, switches to a new host plant (for instance through spill-over from wild reservoirs to cultivated plants) or after its introduction into a new geographic area through global trade (Engering et al. 2013; McCann 2020). The rapid detection of a pathogen in a new environment and/or in a new host plant is therefore crucial to prevent or mitigate disease outbreaks. This is not a trivial task, especially for bacteria with a wide plant host range, many asymptomatic hosts and slow symptom development. Such bacteria can indeed persist undetected for long periods of time in wild reservoirs or even cultivated plants and, once detected, they may already be so well-established that eradication becomes impossible (Soubeyrand et al. 2018; Moralejo et al. 2020). Prime examples for this are the vector-borne xylem-inhabiting bacteria *Xylella fastidiosa* and *X. taiwanensis* (Gammaproteobacteria). *X. fastidiosa* is responsible for numerous diseases in diverse cultivated plants worldwide, notably Pierce’s disease of grapevine, Citrus Variegated Chlorosis, Olive Quick Decline Syndrome and leaf scorches in diverse plants such as coffee and stone fruits (Chatterjee et al. 2008; Rapicavoli et al. 2018). However, the actual plant host range of the bacterium is much larger, since over 700 plant species from 89 families have so far been confirmed as hosts of *X. fastidiosa* in nature or in laboratory experiments (EFSA et al. 2025). It is important to note that many of these plants do not develop diseases, indicating that *X. fastidiosa* associates with most plant species as a commensal rather than as a pathogen (Sicard et al. 2018; Roper et al. 2019). When disease occurs, it is due to the obstruction of xylem vessels by bacterial biofilms and plant tyloses, impairing water transport through the xylem (Newman et al. 2003; Sun et al. 2013).

In addition, *X. fastidiosa* has an important adaptive capacity, as evidenced by its high genetic diversity and frequent recombination events (Jacques et al. 2016; Coletta-Filho et al. 2017; Vanhove et al. 2019). Currently, five subspecies of *X. fastidiosa* have been described (subsp. *fastidiosa*, *multiplex, pauca*) or proposed (subsp. *morus, sandyi*), which evolved in different regions of the Americas. Over the last 150 years, several plant diseases caused by *X. fastidiosa* emerged in the USA and Brazil, for instance after recombination events between subspecies and/or host jumps (Nunney et al. 2012; Nunney et al. 2013; Nunney et al. 2014a; Nunney et al. 2014b; Nunney et al. 2014c; Coletta-Filho et al. 2017; Donegan et al. 2025). Subsequently, *X. fastidiosa* was accidentally introduced to other parts of the world via infected plant material and is now known to be established in Taiwan (Su et al. 2013), China (Guo et al. 2024), the Middle East (Zecharia et al. 2022; Choueiri et al. 2023) and in Southern Europe (France, Italy, Spain and Portugal) (Cariddi et al. 2014; Denancé et al. 2017; Saponari et al. 2019; Landa et al. 2020; Dupas et al. 2023). These introductions caused the emergence of new diseases (Olive Quick Decline Syndrome in Italy as well as Almond Leaf Scorch and Pierce’s disease of grapevine in several countries), accompanied by an expansion of *X. fastidiosa’s* host range to numerous native Mediterranean plants (EFSA et al. 2025).

Hence, introductions of *X. fastidiosa* pose serious threats to both crops and natural ecosystems due to its capacity to establish in new environments. Moreover, new infections can be difficult to detect, due to slow symptom development, symptoms resembling those associated with other stresses (e.g. desiccation) and many asymptomatic hosts. This explains why the presence of the bacterium in some countries remained unnoticed for years prior to its eventual detection (Soubeyrand et al. 2018; Moralejo et al. 2020; Dupas et al. 2023). Consequently, it is conceivable that there are additional unidentified host plants in infected areas as well as yet unrecognised infected areas in other parts of the world. These represent important sources of inoculum from which new outbreaks or further propagations of the bacterium could arise.

In contrast to *X. fastidiosa,* its sister species *X. taiwanensis*, the only additional species described within the genus, has so far only been detected in Taiwan where it causes leaf scorch disease on Asian pear (Su et al. 2016; Weng et al. 2021). Much less is known regarding its genetic diversity and adaptive capacity compared to *X. fastidiosa.* Hence, it is currently unknown whether *X. taiwanensis* is truly restricted to pear and only present in Taiwan or whether the bacterium is also present in other parts of the world, potentially with a larger plant host range.

Since it is impossible to survey all plant species in all environments within the climatic range suitable for *Xylella* survival, the aim of the present work was to identify potential new host plants and infected areas through an in-depth screening of sequence data stored in the NCBI Sequence Read Archive (SRA). As the largest public repository of raw next-generation sequencing data, the SRA provides a unique opportunity to detect the presence of a target organism in sequence data from diverse samples from all over the world. For instance, reads belonging to plant-associated bacteria such as *Xylella* could be present in any plant-derived sequencing datasets that were obtained for completely unrelated research objectives, e.g. to assemble a plant genome. Runs containing reads of the target organism can be identified using the SRA Taxonomic Analysis Tool (STAT), a k-mer-based tool allowing the rapid read-level assignment of taxonomic diversity present in each run (Katz et al. 2021). Reads belonging to the target organism can then be extracted for further analyses. After rigorous identification and elimination of false positives, our SRA data mining revealed potential new host plants and infected areas for both *X. fastidiosa* and *X. taiwanensis*, notably in Africa and Asia. Moreover, we assembled several draft genomes of new strains belonging to *X. fastidiosa* subsp. *multiplex* from North America, *X. fastidiosa* subsp. *pauca* from South America and *X. taiwanensis* from Japan. Furthermore, we obtained the complete circular genome and three draft genomes of a potential new *Xylella* species present in wild rice in Southeast Asia. Taken together, our work presents new insights into the genetic diversity and plant host range of the genus *Xylella* and can help orienting surveillance programmes towards new regions and host plants.

## MATERIALS AND METHODS

### Identification of SRA datasets containing X. fastidiosa sequences

To identify all datasets in which the SRA Taxonomic Analysis Tool (STAT) (Katz et al. 2021) assigned more than 1,000 reads to *X. fastidiosa*, we interrogated the Taxonomy Analysis Table of the nih-sra-datastore on the Google Cloud Platform BigQuery in September 2023 using the following command:

SELECT acc,total_count FROM ‘nih-sra-datastore.sra_tax_analysis_tool.tax_analysis’ WHERE tax_id = 2371 AND total_count > 1000.

The reads of the identified datasets were downloaded in fastq format using the commands *prefetch* and *fasterq-dump* of the SRA Toolkit (https://github.com/ncbi/sra-tools/) or with aria2c via the EBI ftp link.

The presence of *Xylella* spp. sequences in each dataset was verified in two steps (Fig. 1): First, by searching for a set of *Xylella*-specific k-mers located in the 16S rRNA and 23S rRNA genes. This k-mer set had already been established prior to this work, based on all *Xylella* spp. sequences extracted from the SILVA SSU and LSU databases in 2015 (Quast et al. 2013). SkIf2 (Denancé et al. 2019) was used to identify k-mers of variable lengths present in all *Xylella* sequences and absent from all other taxa in the databases. Before using these k-mers in this work, we validated them on the current SILVA 138 databases and verified their presence in all 241 complete or chromosome-level *X. fastidiosa* and *X. taiwanensis* genome assemblies available at NCBI (as of 19/04/2024). Next, we searched for these *Xylella*-specific rRNA k-mers in the sequence datasets identified by STAT using cutadapt (Martin 2011) with the parameters -e 0 -O k-mer length. This approach should detect the presence of *Xylella* spp. in the majority of sequencing projects, notably 16S rRNA amplicon sequencing and whole-genome shotgun sequencing. However, amplicon sequencing projects using a different marker gene (e.g. gyrB) or RNAseq projects involving bacterial ribodepletion would be negative for these ribosomal k-mers.

**Fig. 1.**
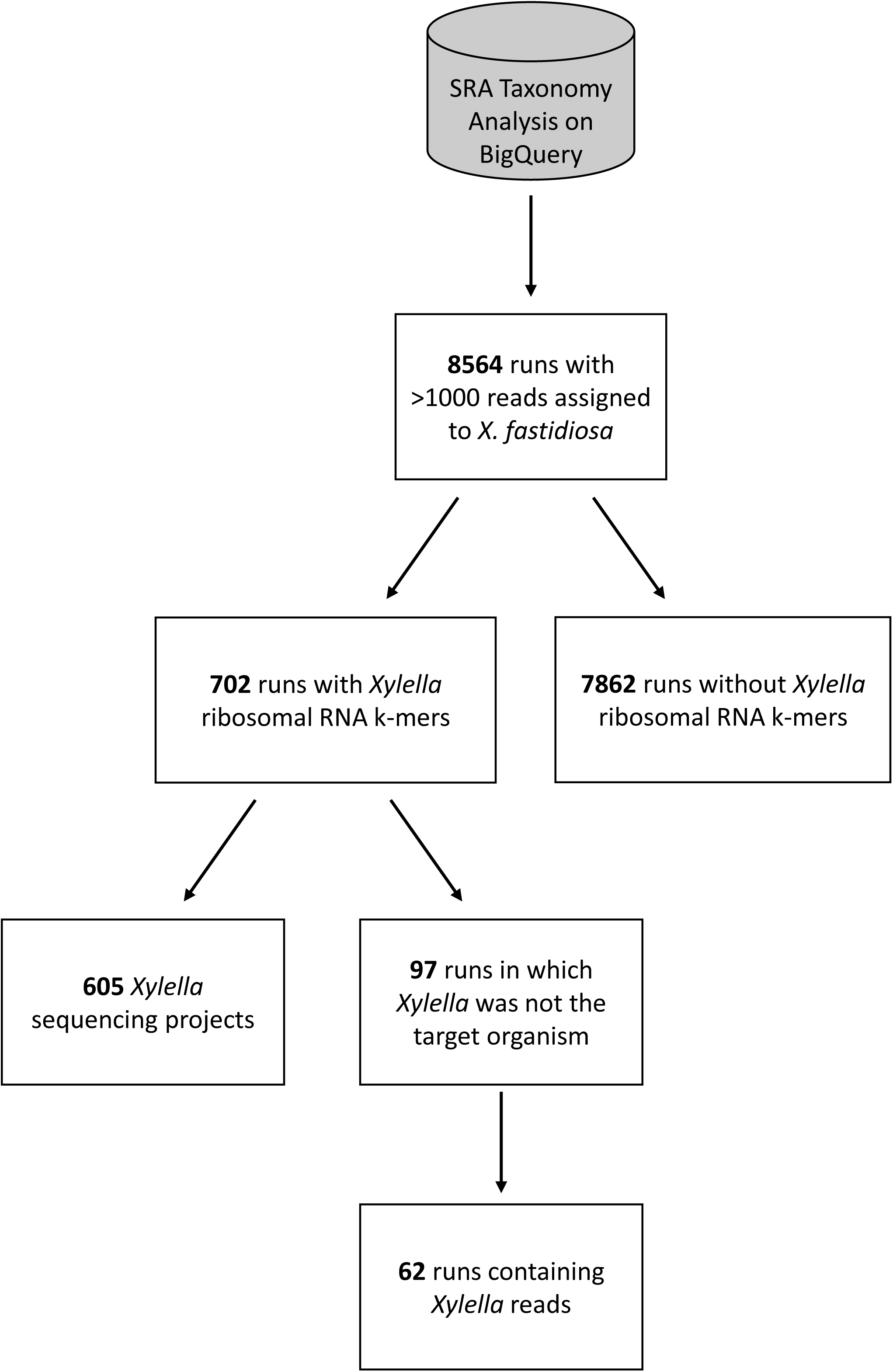
Workflow to validate the presence of *X. fastidiosa* reads in SRA datasets and to eliminate false positives.

Secondly, the datasets that were (a) positive for *Xylella*-specific rRNA k-mers and (b) for which *X. fastidiosa* or *X. taiwanensis* were not the focus of the study were further analysed by mapping the raw reads against a database containing all complete or chromosome-level *X. fastidiosa* and *X. taiwanensis* genome assemblies available at NCBI (as of 19/04/2024) using Bowtie2 v2.5.0 (Langmead and Salzberg 2012). All reads that mapped against this database were extracted for further validation. For 16S amplicon runs, taxonomy was assigned to the mapped reads using the ‘assignTaxonomy’ function of the R package ‘dada2’ (Callahan et al. 2016) against the RDP Training Set 16. Reads assigned to *Xylella* spp. were blasted against the NCBI nt database to verify that the best hit was indeed *Xylella*. If this was not the case, we considered that the data did not contain any *Xylella* spp. reads. For non-amplicon runs, ribosomal rRNA sequences were extracted by mapping against the SILVA database v138.1 and processed in the same way as amplicon sequencing data. Reads that did not map against the SILVA database were blasted directly against the NCBI nt database to verify that the best hit was indeed *X. fastidiosa*.

### Detection of false positives due to contamination of published *X. fastidiosa* genomes

To investigate why STAT reported *X. fastidiosa* reads in whole-genome sequencing (WGS) data that did not contain *X. fastidiosa* according to our rRNA k-mer search, we analysed the reads of 21 such datasets in more detail. First, the raw reads were mapped against a reference database containing all *Xylella* spp. genome assemblies (regardless of assembly quality) available at NCBI as of 20/02/2024 using Bowtie2 v2.5.0 (Langmead and Salzberg 2012). All reads that mapped against the *Xylella* spp. genomes were extracted and *de novo* assembled using Spades v4.0.0 (Prjibelski et al. 2020). The largest contig from each dataset was identified using blast against the NCBI nt database. The sequences of frequent non-*Xylella* blast hits (i.e. plant ribosomal RNAs) were in turn blasted against the *Xylella* spp. genome database to make sure that these contaminating sequences were indeed present in some *X. fastidiosa* genomes. To pinpoint precisely which *X. fastidiosa* genomes and contigs contain these plant-derived contaminants, we used Kmer-db v1.9.2 (Deorowicz et al. 2019) to build a database of all k-mers of size 22 contained in all published *X. fastidiosa* genome assemblies. Subsequently, the command kmer-db new2all was used to investigate the distribution of each k-mer across all *X. fastidiosa* genomes. The rationale behind this analysis was that if the majority of k-mers occurring on the same contig were present only in a single genome, then this contig would be a likely candidate for a contamination.

### Subspecies identification in X. fastidiosa-positive runs

All publicly available *X. fastidiosa* and *X. taiwanensis* genomes were grouped according to the percentage of shared K-mers using the tool KI-S (Briand et al. 2021). SkIf2 (Denancé et al. 2019) was then used to determine specific k-mers for each *X. fastidiosa* subspecies and for *X. taiwanensis* and to count these k-mers in each non-amplicon dataset containing *Xylella* reads. This approach was chosen to increase the chances to assign reads to species and subspecies even for datasets with very low abundance of *Xylella* reads where it may not be possible to assemble the complete sequences of the seven housekeeping genes used in the *Xylella* MLST scheme (Yuan et al. 2010). After the detection of a new *Xylella* species from several wild rice datasets (see Results), the diagnostic k-mers were updated to distinguish also between *X. taiwanensis* and the new *Xylella* species.

### Xylella sp. genome assemblies

*Xylella* sp. draft genomes were assembled from 10 datasets with high numbers of reads assigned to *Xylella* via mapping against the *Xylella* spp. whole-genome reference database mentioned above. Illumina reads were assembled using Spades v4.0.0 (Prjibelski et al. 2020) with the parameters --cov-cutoff auto --isolate and Unicycler v0.4.9 (Wick et al. 2017) followed by one iteration of Redundans v0.14 (Pryszcz and Gabaldon 2016) for scaffolding and gap-closing. Small contigs <500 bp were discarded. Oxford Nanopore reads from SRR12451683 (WGS of wild rice) were assembled using Flye v2.9.1 (Kolmogorov et al. 2020) and polished with three iterations of Polca (Zimin and Salzberg 2020), using Illumina reads from the same plant sample (SRR13453590) (Shang et al. 2022). As this assembly was quite different from known *Xylella* strains (see Results), it was added to the *Xylella* spp. genome database to extract more *Xylella* reads from other SRA runs from wild rice. Genome completeness and contamination was assessed using CheckM v1.2.2 (Parks et al. 2015) and BUSCO v5.4.4 (Manni et al. 2021) with the database gammaproteobacteria_odb10. Assemblies were annotated using Bakta v1.5.1 (Schwengers et al. 2021) and MLST sequence typing was done using the *Xylella* PubMLST database (https://pubmlst.org/organisms/xylella-fastidiosa). Average nucleotide identity (ANI) was calculated using the EZBioCloud ANI calculator (https://www.ezbiocloud.net/tools/ani).

### Phylogenomic analysis

A core-genome phylogenomic analysis was performed using Orthofinder v2.5.4 (Emms and Kelly 2019) to identify all single-copy orthologous genes shared between the new *Xylella* sp. draft genomes, 79 genomes covering the genetic diversity of *X. fastidiosa* and *X. taiwanensis* and three genomes of *Xanthomonas albilineans* as outgroup. Only complete genomes were included in this analysis, unless when no complete genome assembly is currently available (e.g. for *X. fastidiosa* sequence types 8, 9, 87). The amino acid sequences of each conserved gene were aligned using Muscle v5.2 (Edgar 2004) and the alignments were concatenated into a partitioned supermatrix using the script geneStitcher.py (https://github.com/ballesterus/Utensils/blob/master/geneStitcher.py). IQ-TREE v1.6.12 (Minh et al. 2020) was used to predict the optimal amino acid substitution model for each gene partition (Chernomor et al. 2016; Kalyaanamoorthy et al. 2017) and to produce a Maximum Likelihood phylogenetic tree with 1000 bootstrap iterations. The tree was visualized in FigTree v1.4.4 (https://github.com/rambaut/figtree).

### KEGG pathway and secretion system analysis

KEGG pathway annotations were obtained using BlastKOALA v3.1 (Kanehisa et al. 2016) for the four genome assemblies of the new *Xylella* species from wild rice as well as for seven reference strains from *X. fastidiosa* and *X. taiwanensis* (Temecula, WM1-1, AlmaEM3, M12, 9a5c, DeDonno and PLS229). Pathway completeness was calculated using KEGG Decoder v1.3 (Graham et al. 2018). MacSyFinder 2.0 implemented in the tool TXSScan (Abby et al. 2016) was used to identify bacterial secretion systems in 78 complete *Xylella* spp. genomes (71 from all *X. fastidiosa* subspecies, 6 from *X. taiwanensis* and our circular genome assembled from wild rice). Completeness of KEGG pathways and bacterial secretion systems were visualized in heatmaps using the R package “pheatmap” in R v4.4.3 (R Core Team 2021).

## RESULTS

### Overestimation of *X. fastidiosa* sequences in SRA datasets due to contaminated genome sequences

Our SRA query identified 8,564 datasets in which the SRA Taxonomic Analysis Tool (STAT) (Katz et al. 2021) assigned more than 1,000 reads to *X. fastidiosa* (Supplementary Table S1). SkIf2 (Denancé et al. 2019) initially identified 20 *Xylella*-specific rRNA k-mers of variable lengths (22-43 nucleotides) in the sequences assigned to *Xylella* spp. in the SILVA SSU and LSU databases (Supplementary Table S2). However, in the course of this work we progressively discarded 12 of these k-mers since they were found in numerous SRA datasets that did not in fact contain any *Xylella* reads upon closer inspection via read mapping. Therefore, we only searched for the remaining eight k-mers (K1, K2, K4, K5, K6, K7, K9 and K20) in the raw reads of the 8,564 SRA datasets and applied a minimum threshold of at least ten counts (considering that the same k-mer can be counted multiple times) to avoid artefacts. This resulted in only 702 datasets in which at least ten counts of the eight *Xylella*-specific rRNA k-mers were detected (Fig. 1). Hence, 91.80% of the datasets reported positive for *X. fastidiosa* by STAT (7,862 out of 8,564) did not contain any *X. fastidiosa* ribosomal RNA sequences.

To investigate why the Taxonomic Analysis Tool reported *X. fastidiosa* reads in so many datasets that did not contain *X. fastidiosa* according to our rRNA k-mer search, we analysed a subset of 21 WGS datasets that were negative in our k-mer analysis in more detail. All of these datasets contained indeed reads that mapped against a database containing all published *X. fastidiosa* genome assemblies. The mapped reads were assembled into contigs and the largest contig from each dataset was blasted against the NCBI nt database. Surprisingly, the majority (14/21) of these contigs returned plant ribosomal RNAs as best blast hits, whereas the remaining seven corresponded to a miscRNA of *Arabidopsis thaliana*, mouse rRNAs, rat mitochondrion and phages (Supplementary Table S3). Blasting these sequences back against the *X. fastidiosa* genome database revealed that these contaminating sequences were indeed present in *X. fastidiosa* genomes, often on a single small contig of a single genome (Supplementary Table S3). To better identify the contigs containing such contaminants in the published *X. fastidiosa* genomes, we used Kmer-db to determine all possible 22-mers across all contigs of all published genomes and then analysed the distribution of these k-mers. We reasoned that if a k-mer is only found on a single contig in one genome, there is a high probability that this contig is a contamination. We identified 871 such isolated contigs (Supplementary Table S4). These were present in 26 genomes and were generally very small (209-16,556 bp). Blasting these contigs against the NCBI nt database returned diverse other bacteria as best hits for 448 contigs (51%), 239 contigs (27%) had no hit, 48 contigs (5.5%) corresponded to animal (mostly rat) sequences, 44 contigs (5%) to plant sequences, 40 contigs (4.6%) to mitochondria of animals and plants, 33 contigs (3.8%) to chloroplasts, 13 contigs (1.5%) to *X. fastidiosa* and 6 contigs (0.7%) to animal or plant rRNAs (Supplementary Table S4). It is important to note that the 26 genomes in which these contigs were identified are all draft genomes with hundreds of contigs while no singleton k-mers were found in any complete or chromosome level genome assemblies. Hence, it is likely that most of these contigs are contaminating sequences, potentially derived from cross-contamination or index hopping when several projects are sequenced together in the same run.

### Identification of SRA datasets containing *Xylella* spp. sequences

Of the 702 datasets positive for *Xylella*-specific rRNA k-mers, 605 corresponded to studies investigating *X. fastidiosa,* e.g. genome sequencing projects or amplicon sequencing of *X. fastidiosa* host plants (Fig. 1). This left us with 97 datasets for which *X. fastidiosa* was not the focus of the study and reads affiliated to *X. fastidiosa* or *X. taiwanensis* were identified in 62 of these datasets via mapping or amplicon analysis (Table 1, Supplementary Table S5). In the other cases, the reads that mapped against the *Xylella* spp. reference genomes did not return *Xylella* as best hit by blast analysis against the NCBI nt database. Therefore, the presence of *Xylella* in those datasets could not be confirmed.

**Table 1.** Summary of the 62 SRA datasets containing *Xylella* spp. reads in new host plants and localities. Species and subspecies identification is based on k-mer counts reported in Supplementary Table S5. Datasets from which complete or draft genomes could be assembled are highlighted in bold.

| Technology | Sample type | Host species | Tissue | Location | *Xylella* species/subspecies | # Datasets | References |
| --- | --- | --- | --- | --- | --- | --- | --- |
| RADseq | Plant | *Alloteropsis angusta* | Leaf | Zambia | *Xylella* sp. | 2 | Curran et al. 2022 |
| Amplicon | Plant | *Cannabis sativa* | Leaf | USA | ND | 5 | Ahmad et al. 2024 |
| Amplicon | Plant | *Cannabis sativa* | Root | USA | ND | 2 | Ahmad et al. 2024 |
| Amplicon | Plant | *Cannabis sativa* | Stem | USA | ND | 7 | Ahmad et al. 2024 |
| RNAseq | Plant | *Coffea arabica* | Leaves | Brazil | *X. fastidiosa* subsp. *pauca* | 7 |  |
| **WGS** | **Plant** | ***Coffea arabica*** | **Leaves** | **NA** | ***X. fastidiosa* subsp. *pauca*** | **2** |  |
| **WGS** | **Plant** | ***Coffea eugenoides*** | **Leaves** | **Colombia** | ***X. fastidiosa* subsp. *pauca*** | **1** |  |
| **WGS** | **Plant** | ***Isodon japonicus*** | **Leaf** | **Japan** | ***X. taiwanensis*** | **1** | Chen et al. 2022 |
| **WGS** | **Plant** | ***Lavandula stoechas*** | **Leaf** | **USA** | ***X. fastidiosa* subsp. *multiplex*** | **1** |  |
| WGS | Plant | *Lavandula x intermedia* | Leaf | USA | *X. fastidiosa* subsp. *multiplex* | 1 |  |
| Amplicon | Plant | *Musa acuminata* | Rhizosphere | Nicaragua | ND | 2 | Köberl et al. 2015, 2017 |
| **WGS** | **Plant** | ***Oryza rufipogon*** | **Leaf** | **China** | ***X. rufipogonis*** | **2** | Shang et al. 2022 |
| **WGS** | **Plant** | ***Oryza rufipogon*** | **Leaf** | **Myanmar** | ***X. rufipogonis*** | **2** | Shang et al. 2022 |
| Amplicon | Plant | *Populus deltoides* | Leaf | USA | ND | 2 | Argiroff et al. 2024 |
| Amplicon | Plant | *Populus trichocarpa* | Leaf | USA | ND | 10 | Argiroff et al. 2024 |
| **WGS** | **Plant** | ***Populus trichocarpa*** | **NA** | **USA** | ***X. fastidiosa* subsp. *multiplex*** | **1** |  |
| Amplicon | Plant | *Vitis vinifera* | Cane | USA | ND | 3 | Deyett et al. 2017 |
| RNAseq | Fly | *Xiphidiella anorubra* | Whole body | Namibia | *X. fastidiosa* subsp. *multiplex* | 1 | Yan et al. 2021 |
| RNAseq | Millipede | *Chondrodesmus cairoensis* | Head | Costa Rica | *X. fastidiosa* subsp. *fastidiosa* | 1 | Rodriguez et al. 2018 |
| RNAseq | Millipede | *Orthoporus* sp. | Head | Costa Rica | *X. fastidiosa* subsp. *fastidiosa* | 1 | Rodriguez et al. 2018 |
| Amplicon | Human | *Homo sapiens* | Gut | Italy | ND | 1 | Russo et al. 2021, 2022 |
| RNAseq | Human | *Homo sapiens* | Granulation tissue | USA | *X. fastidiosa* subsp. *multiplex* | 2 |  |
| WGS | Human | *Homo sapiens* | Faeces | USA | *X. fastidiosa* subsp. *pauca* | 1 |  |
| WGS | Bacteria | *Clavibacter lycopersici* | Bacterial culture | Iran | *X. fastidiosa* subsp. *multiplex* | 1 |  |
| **WGS** | **Bacteria** | **Mock community** | **Bacterial culture** | **Belgium** | ***X. fastidiosa* subsp. *fastidiosa*** | **1** | Goussarov et al. 2024 |
| WGS | Bacteria | Mock community | Bacterial culture | Switzerland | *X. fastidiosa* subsp. *pauca* | 2 |  |

32 datasets contained 16S rRNA amplicon sequencing data from five different plant species (*Cannabis sativa, Musa accuminata* (banana), *Populus deltoides, Populus trichocarpa, Vitis vinifera*) and, surprisingly, also a human gut metagenome (Table 1). While the presence of *Xylella* as an abundant taxon had already been described in *C. sativa* and grapevine (Deyett et al. 2017; Ahmad et al. 2024), it had not been reported previously from the microbiomes of *M. accuminata*, *Populus* spp. and the human gut (Köberl et al. 2015; Köberl et al. 2017; Russo et al. 2021; Russo et al. 2022; Argiroff et al. 2024), likely because it was not among the highly abundant taxa or not a taxon of interest for the respective studies. The *Xylella* species cannot be determined from short amplicon sequences using our specific k-mers. However, considering that all plant samples came from the USA except for banana, it is likely that these plants were infected with *X. fastidiosa*. The sequenced bananas were sampled in Nicaragua, a country in which *X. fastidiosa* has not yet been reported, despite its presence in neighbouring Costa Rica (https://gd.eppo.int/taxon/XYLEFA/distribution). Since *Xylella* was detected in the rhizosphere soil and not in the phyllosphere, it is possible that the bacterium did not infect banana itself, but instead a different crop planted in association with banana (coffee in this case) (Köberl et al. 2015).

The remaining 30 datasets corresponded to RADseq, RNAseq and whole-genome sequencing (WGS) data from diverse plant species sampled in Africa, Asia and the Americas as well as several invertebrates (millipedes and a flesh fly), human tissues or faeces collected in medical facilities and several bacterial mock communities (Table 1). One of the latter validated our approach, considering that the mock community sequenced in run ERR5321934 contained the *X. fastidiosa* subsp. *fastidiosa* strain LMG 17159 (synonym of ATCC 35879, CFBP 7970, DSM 10026, ICMP 15197) (Goussarov et al. 2024). We assembled a good-quality draft genome from the data that shared 99.96% ANI with the published genome of ATCC 35879 and clustered together with several *X. fastidiosa* subsp. *fastidiosa* sequence type (ST) 2 strains in the phylogenomic analysis (Table 2, Fig. 2). Astonishingly, *Xylella* reads were also detected in the WGS data from a pure culture of *Clavibacter lycopersici* CFBP 8616. Since this sequencing data had been produced in our own lab, we could trace back that a *Xylella* strain had been multiplexed with *C. lycopersici* in the same run. Hence, the presence of *Xylella* reads among the *C. lycopersici* raw data can be explained by cross-contamination or index hopping during the library preparation and sequencing and does not indicate a contaminated *C. lycopersici* culture or incorrect genome assembly.

**Fig. 2.**
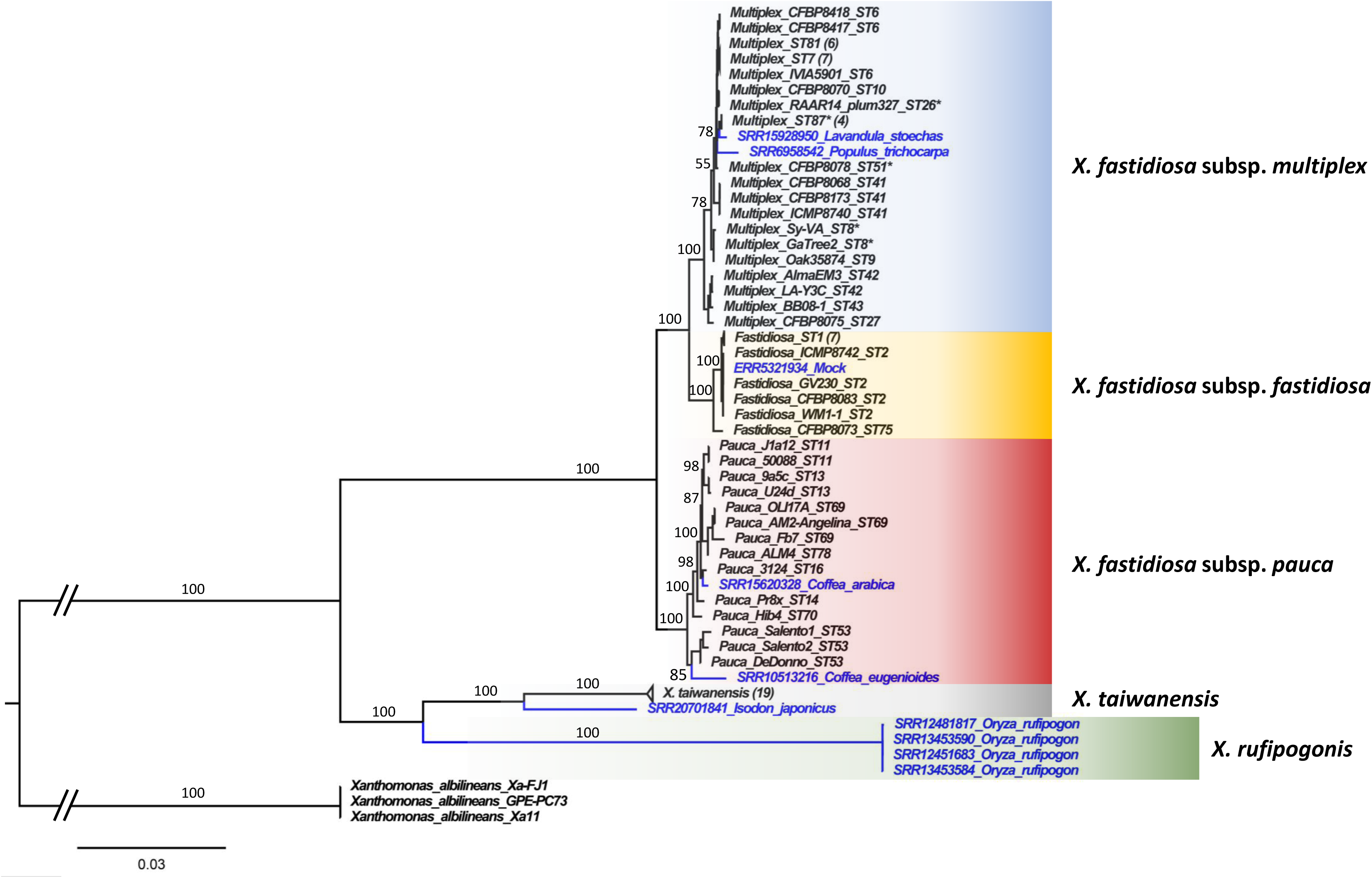
Phylogenomic analysis revealing numerous new strains and a potential new *Xylella* species associated with rice. Maximum-likelihood phylogenetic tree based on 146 single-copy protein-coding genes shared between 89 *Xylella* spp. strains and 3 *Xanthomonas albilineans* strains as outgroup. *Xylella* genomes assembled from SRA datasets are indicated in blue. Numbers in parentheses indicate the number of genomes included in collapsed clades. Asterisks indicate incomplete draft genomes. Branch support is based on 1000 bootstrap iterations and is provided only for major branches in the interest of readability.

**Table 2.**
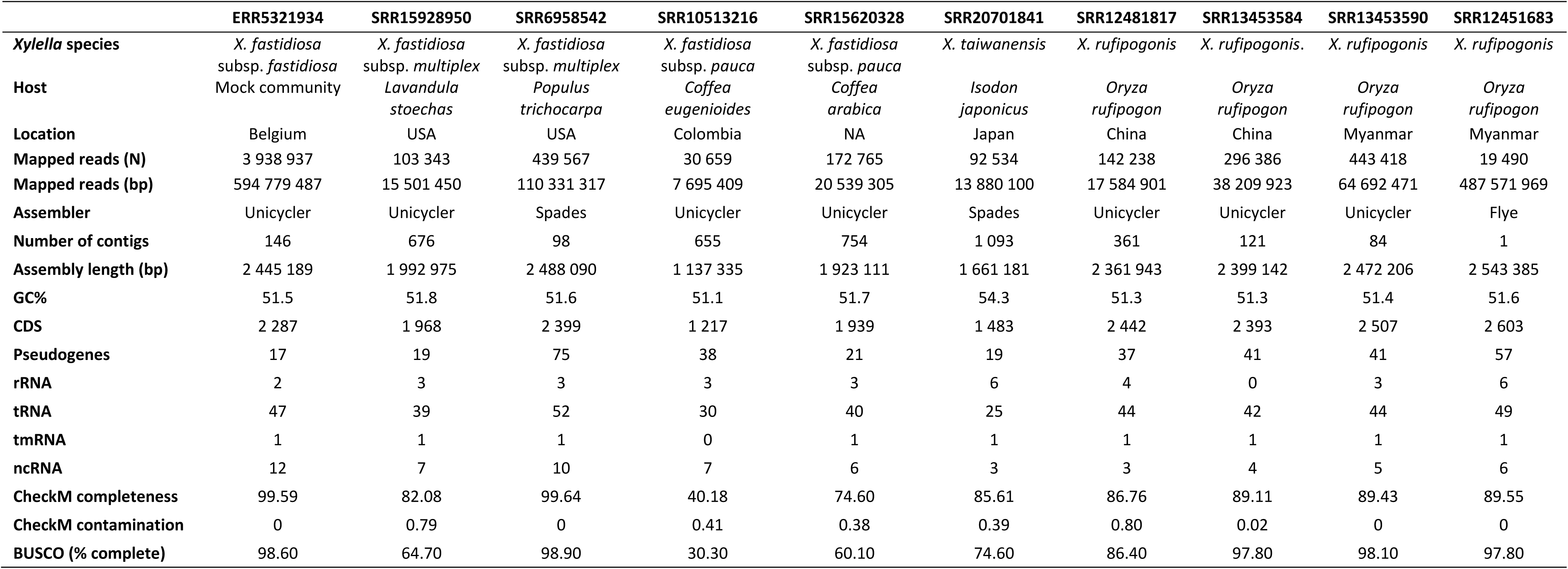
Summary of the *Xylella* spp. genomes assembled in this study.

For the RADseq, RNAseq and WGS datasets, the *Xylella* species and *X. fastidiosa* subspecies were determined based on specific k-mers (Supplementary Table S5), suggesting several potential new host plants and a larger geographic range for both *Xylella* species (Table 1). Hence, *X. fastidiosa* subsp. *pauca* was detected in several datasets from coffee plants in Brazil as well as Colombia. While this subspecies is known as the causal agent of coffee leaf scorch in Brazil (Nunney et al. 2012; Coletta-Filho et al. 2017), the presence of *X. fastidiosa* in Colombia was declared for the first time in March 2025 (https://gd.eppo.int/taxon/XYLEFA/distribution/CO). The subspecies *multiplex* was detected in two species of lavender from the USA, already known as host plants of this subspecies (EFSA et al. 2025), as well as for the first time in black cottonwood (*Populus Trichocarpa)* (in line with the amplicon sequence data mentioned above) and in a fly from Namibia. While it is unlikely that *X. fastidiosa* was alive in a fly that does not possess the appropriate mouthparts to pump xylem sap and to transfer vascular bacteria between plants, it is conceivable that the bacterium was present in the environment (e.g. associated with particular plants) and that its DNA was present in or on the fly at the time of sampling. The possibility that these reads result from a cross-contamination between samples during library preparation or sequencing cannot be entirely excluded but seems unlikely due to the relatively high number of *Xylella* reads in this dataset (Supplementary Table S5). Nonetheless, the amount of *Xylella* reads was insufficient for genome assembly, which produced only numerous short contigs for a total length of 264 Kbp. In contrast, the detection of *X. fastidiosa* subsp. *fastidiosa* in the heads of two millipede species from Costa Rica is less surprising, considering that this subspecies is known to occur in Costa Rica and the majority of millipedes are detritivores feeding on decaying plant material. The presence of *X. fastidiosa* in several human tissue samples cannot be explained at present. However, it is worth noting that these samples originated exclusively from two countries in which *X. fastidiosa* is well-established, i.e. Italy and the USA. Therefore, it cannot be excluded that these *Xylella* reads might also be the result of cross-contaminations or index hopping during sample multiplexing, as outlined above for the *Xylella* reads detected in a *Clavibacter* dataset.

K-mers specific for *X. taiwanensis* were initially detected in three plant species collected in four countries: The African grass *Alloteropsis angusta* from Zambia, wild rice *Oryza rufipogon* from Myanmar and China and *Isodon japonicus* (Lamiaceae family) from a botanical garden in Japan. However, in a second analysis using revised k-mers able to distinguish between *X. taiwanensis* and a new *Xylella* species associated with wild rice (see the phylogenomics analysis described below), only the reads from *I. japonicus* could be firmly assigned to *X. taiwanensis* (Table 1, Supplementary Table S5). It is noteworthy that the *A. angusta* dataset presented similar k-mer counts for *X. taiwanensis* and the new *Xylella* sp., but this result may not be representative due to the overall low number of *Xylella* reads retrieved from these datasets.

### Genome assemblies reveal numerous new strains and a new *Xylella* species associated with wild rice

Although most SRA datasets contained low proportions of *Xylella* reads based on read mapping (Supplementary Table S5), ten draft genomes could be assembled (Table 2). Most genomes are quite fragmented and some are incomplete, with assembly sizes ranging from 1.13 Mbp in 655 contigs to 2.48 Mbp in 98 contigs (Table 2). Nonetheless, we also obtained a complete circular genome of 2.54 Mbp from Oxford Nanopore Promethion data from wild rice (SRR12451683). Initially, this assembly presented many frameshifts and an inflated number of genes compared to typical *Xylella* genomes due to the lower quality of the Nanopore reads. Many of these could be corrected by polishing with Illumina reads from the same plant sample (SRR13453590), achieving similar gene numbers and CheckM completeness score as the more fragmented assembly obtained from the Illumina data from SRR13453590 alone. The most complete genomes in terms of CheckM marker genes are a *X. fastidiosa* subsp. *multiplex* strain from black cottonwood (*Populus trichocarpa*, SRR6958542) and the aforementioned *X. fastidiosa* subsp. *fastidiosa* strain from a mock community (ERR5321934). Both assemblies are >99.5% complete with no contamination (Table 2).

A phylogenomic analysis based on 146 single-copy protein-coding genes shared between 89 *Xylella* spp. genomes and three genomes from *Xanthomonas albilineans* as outgroup revealed an unexpected genetic diversity for the nine draft genomes obtained from the above-mentioned plant samples (Fig. 2): Two of them represent different strains of *X. fastidiosa* subsp. *multiplex* from Spanish lavender (*Lavandula stoechas*) and black cottonwood (*P. trichocarpa*) in the USA. The strain from lavender was most closely-related to ST87 strains from diverse ornamental plants (including *Lavandula* spp.) in Italy (Saponari et al. 2019) and five of the seven MLST genes perfectly matched the ST87 alleles. The two other genes (*leuA* and *nuoL*) were incomplete, precluding a complete ST determination for this strain. In contrast, the strain from *P. trichocarpa* clearly represents a new sequence type: It was placed as an isolated but strongly supported (78% bootstrap support (BS)) branch in the phylogenetic tree and three of the seven MLST genes (*gltT, holC* and *nuoL*) represented new alleles.

In addition, two different strains of *X. fastidiosa* subsp. *pauca* were assembled from *Coffea* spp. in South America (Fig. 2). Their sequence types could not be determined since several MLST genes were missing from both assemblies. Nonetheless, the strain from *C. arabica* was closely-related to ST16 strain 3124 (BS: 98), also isolated from a coffee plant in Brazil. ST16 strains have so far only been observed on coffee (and to a lesser extent olive trees) in Brazil (Coletta-Filho et al. 2017). The strain from *C. eugenioides* on the other hand was most closely related to ST53 strains isolated from olive trees in Italy (BS: 85), which belong to a clonal complex of strains infecting coffee and oleander in Costa Rica (Marcelletti and Scortichini 2016; Giampetruzzi et al. 2017). Hence, our data indicates that related strains are also present in coffee in Colombia.

The remaining five strains did not belong to the species *X. fastidiosa.* Instead, the strain assembled from *Isodon japonicus* in Japan formed a sister branch to *X. taiwanensis* strains from Taiwan (BS: 100, Fig. 2) and might still belong to the same species, as it has 94.03% average nucleotide identity (ANI) with *X. taiwanensis* (strain PLS229) and only 86.0% ANI with *X. fastidiosa* (strain Temecula). The generally accepted cut-off for species delimitation is 95% ANI (Olm et al. 2020) but since our genome assembly is incomplete (1.66 Mbp total assembly length), we consider it premature to consider this strain a different species from *X. taiwanensis.* Finally, we assembled four highly similar *Xylella* genomes from wild rice (*Oryza rufipogon*) collected in China and Myanmar. Interestingly, these strains formed a sister clade to *X. taiwanensis* (BS: 100, Fig. 2) and had only 83.73-83.98% ANI with this species (strain PLS229) and even less (80.92-81.02%) with *X. fastidiosa* (strain Temecula). Hence, the strains assembled from wild rice represent a new species within the genus *Xylella*, for which we propose the name *Xylella rufipogonis*.

### X. rufipogonis has a unique repertoire of secretion systems

To investigate whether *X. rufipogonis* possesses specific functions that might be involved in the interaction with its host plant, we first compared KEGG pathway completeness in the four *X. rufipogonis* genomes to seven reference strains from *X. fastidiosa* and *X. taiwanensis* (Temecula, WM1-1, AlmaEM3, M12, 9a5c, DeDonno and PLS229). This analysis showed that *X. rufipogonis* was very similar to the other strains, with only four KEGG categories being differentially represented (Supplementary Fig. S1): 1) Cellulases, which were absent in *X. rufipogonis* but present in all reference strains, 2) chemotaxis, which was incomplete in *X. rufipogonis* but entirely absent from the reference strains, 3) type I secretion (T1SS), which was less complete in *X. rufipogonis* compared to the reference strains, and 4) type IV secretion (T4SS), which was more complete in *X. rufipogonis* compared to most reference strains except the strain 9a5c. This makes sense because the latter possesses conjugative transfer operons (*tra-trb*) on both the main chromosome and its plasmid (Pierry et al. 2020).

Considering the importance of bacterial secretion systems for the interaction with other bacteria, host cells or the extracellular environment, we performed a finer screening of 78 complete *Xylella* spp. genomes (71 from all *X. fastidiosa* subspecies, 6 from *X. taiwanensis* and our circular genome assembled from wild rice) for all currently defined secretion systems using TXSScan (Abby et al. 2016). Only secretion systems present on the main chromosomes were considered, to avoid biases due to the T4SS conjugative systems present on plasmids in 50% of the strains. This screening revealed that all 78 strains encode a complete T2SS, a type 4 pilus and two to three T5SS autotransporters (Fig. 3). This is not surprising, since the T2SS is required for the successful colonization of the xylem (Ingel et al. 2023) and the type 4 pilus is known to be involved in upstream mobility (Meng et al. 2005). In contrast, T1SS and T4SS were rare, the former being present only in *X. rufipogonis, X. taiwanensis* and a single *X. fastidiosa* strain (strain Hib4 from the subsp. *pauca*) (Fig. 3). T4SS were not detected in any *X. taiwanensis* and *X. fastidiosa* subsp. *multiplex* strains, but conjugative T4SS systems (mainly *trb* operons) were found in *X. rufipogonis,* the *X. fastidiosa* subsp. *fastidiosa* strain Stag’s Leap, two *X. fastidiosa* subsp. *sandyi* strains (Ann-1, OC8), all five *X. fastidiosa* subsp. *morus* strains and four *X. fastidiosa* subsp. *pauca* strains from Brazil (9a5c, Hib4, J1a12, U24D) (Fig. 3). The T4SS operon of *X. rufipogonis* was the most complete in comparison to the other strains. Hence, *X. rufipogonis* possesses a unique and potentially more complete repertoire of secretion systems than all previously sequenced *Xylella* spp. strains.

**Fig. 3.**
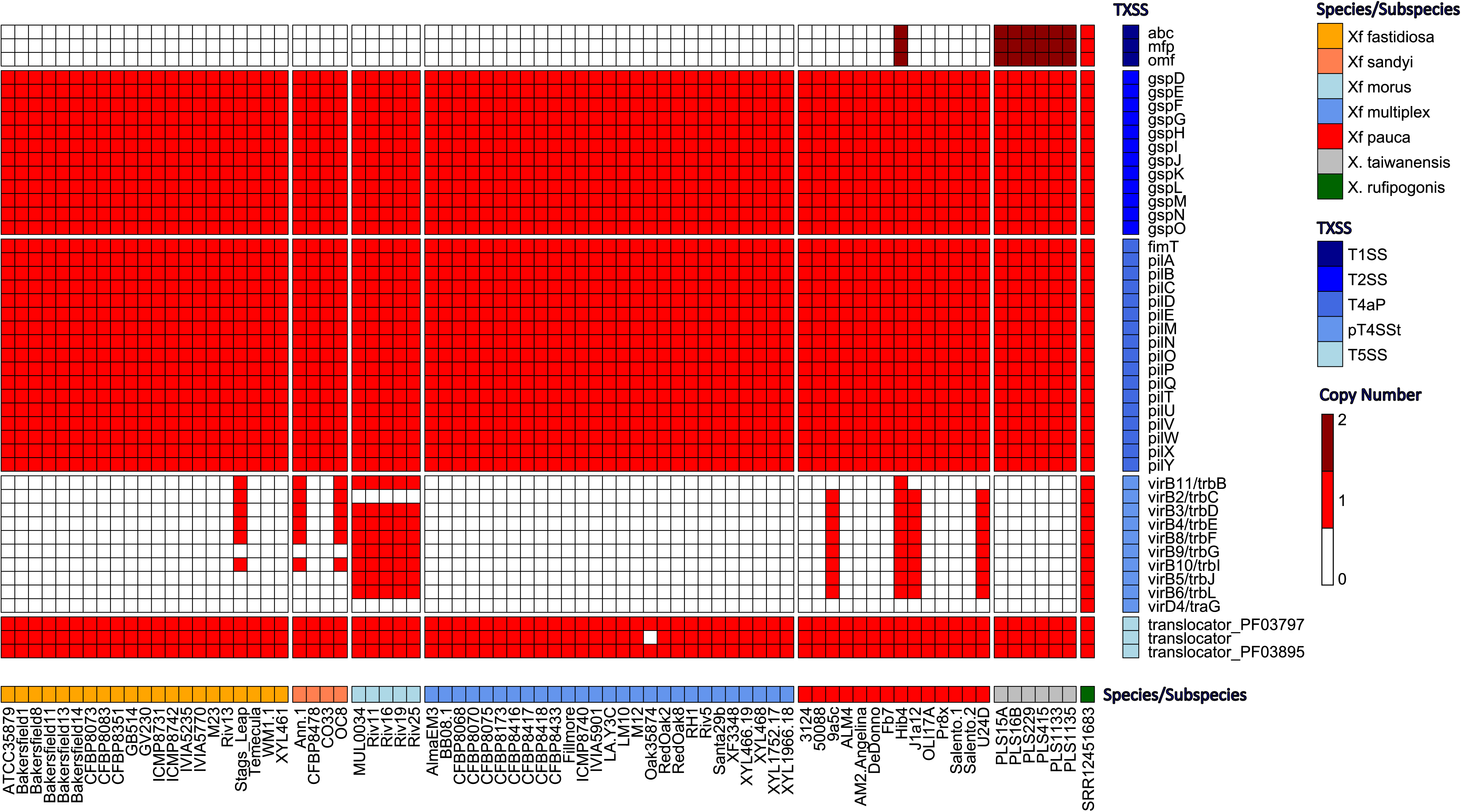
*X. rufipogonis* possesses a unique repertoire of bacterial secretion systems. Heatmap showing the distribution of bacterial secretion systems across 78 complete *Xylella* spp. genomes. Columns are organised by *Xylella* species/subspecies, ordered based on their phylogenetic relationships as in (Jacques et al. 2016; Denancé et al. 2019). Rows are organised by secretion system. Only secretion systems encoded on the main chromosome were included.

## DISCUSSION

In this work, we searched for *Xylella* spp. in diverse plant and animal samples from around the world through data mining of raw sequence reads deposited in the SRA. Considering the huge amount of sequence data currently available, we used the SRA Taxonomic Analysis Tool (STAT) (Katz et al. 2021) to identify only those datasets that might contain *Xylella* reads. Surprisingly, the presence of 16S rRNA gene sequences belonging to *Xylella* spp. could only be confirmed in less than 10% of the >8,500 datasets returned by STAT. It cannot be excluded that some datasets containing *Xylella* reads were missed by our 16S rRNA gene search if ribosomal genes were excluded during library preparation (e.g. amplicon sequencing based on non-ribosomal marker genes or RNAseq projects involving bacterial ribodepletion). However, it seems unlikely that this could be the sole explanation for this discrepancy since these sequencing methods are not the most widely used. Instead, we suspect that a non-negligible fraction of the STAT results represents false positives, especially considering the presence of contigs containing animal or plant sequences that we detected in a few published *X. fastidiosa* genomes. These could indeed explain the erroneous detection of *X. fastidiosa* reads by STAT, considering that the tool classifies query reads based on a reference k-mer database built from published genomes (Katz et al. 2021). Therefore, the presence of contaminating sequences in a published genome assembly can lead to k-mers being wrongly associated with a given organism in the NCBI taxonomy, impacting all downstream analyses such as the taxonomic assignment of reads. Hence, we hope that the increased quality of *Xylella* genomes that we have seen in recent years will ultimately lead to an optimisation of the diagnostic k-mers used by STAT or other tools. In the meantime, careful verifications like those applied in this work are required before concluding that reads of the target organism are indeed present in the datasets retained for further analyses. Another complication may arise from cross-contaminations between samples from different projects that are multiplexed in the same sequencing run. This can result in some reads being assigned to the wrong biological sample during demultiplexing, e.g. via index-hopping. The detection of *Xylella* reads in the raw WGS data from a *Clavibacter* strain multiplexed with a *Xylella* strain on the same run illustrates this risk. Hence, it cannot be excluded that we missed additional datasets and a few of the 62 datasets in which we could validate the presence of *Xylella* reads may not actually be biologically meaningful (e.g. bacterial mock communities and data from humans).

Nonetheless, most of the identified datasets from diverse plants and insects produced exciting new insights into the genetic diversity, host range and distribution areas of several *Xylella* species. Many of these findings were further supported by the assembly of eight draft genomes and one circular genome from different plant species sampled around the world. For instance, we identified two new strains of *X. fastidiosa* subsp. *multiplex* in the USA, one from Spanish lavender (*Lavandula* stoechas) and the other from black cottonwood (*Populus trichocarpa)*, a species not previously reported as a host plant of *X. fastidiosa* (EFSA et al. 2025). The strain from *L. stoechas* is closely related to ST87 strains so far only detected in Italy (in various plants including *Lavandula* spp. (Saponari et al. 2019)) and might even belong to the same sequence type, indicating another probable introduction event from the USA to Europe. In contrast, the strain from *P. trichocarpa* represents a new sequence type, thus increasing our knowledge of the genetic diversity of the subspecies *multiplex* in deciduous trees native to North America. In addition, we identified two new strains of *X. fastidiosa* subsp. *pauca* from *Coffea* spp. in Brazil and Colombia. The strain from *C. eugenioides* from Colombia is of particular interest, as it represents a new genetic lineage most closely-related to a clade of ST53 strains known to infect coffee and oleander in Costa Rica and olive trees in Italy (Marcelletti and Scortichini 2016; Giampetruzzi et al. 2017).

Integrating both genomic and amplicon datasets, our results indicate the presence of *X. fastidiosa* in two additional countries: Nicaragua (16S rRNA amplicons in the rhizosphere of banana planted in association with coffee (Köberl et al. 2015)) and Namibia (*X. fastidiosa* subsp. *multiplex* reads in a fly transcriptome). The latter may be the most surprising finding since *Xylella* has so far not been detected in Africa. Unfortunately, the amount of potential *Xylella* reads from this dataset was insufficient for draft genome assembly, suggesting that the fly is not a true host of *X. fastidiosa* but simply carried some bacterial cells or DNA in or on its body at the time of sampling. Interestingly, this was not the only dataset from Africa - reads assigned to *Xylella* spp. based on specific k-mers were also detected in RADseq data from the perennial African grass *Alloteropsis angusta* sampled in Miombo woodlands in Zambia (Curran et al. 2022). Unfortunately, it is not possible to assemble whole genomes from RADseq data but we obtained a few short contigs representing different genes, making cross-contamination from amplicon sequencing unlikely. In contrast, the partial genome of a new strain likely belonging to *X. taiwanensis* could be assembled from *Isodon japonicus* (Lamiaceae family) from a botanical garden in Japan and four genomes (one of them circularized) were assembled from leaves of wild rice (*Oryza rufipogon*) sampled in China and Myanmar (Shang et al. 2022). The genomes from both countries were highly similar and both ANI and phylogenomic analyses support the conclusion that these strains represent a new *Xylella* species, for which we propose the name *X. rufipogonis.* The circularized genome of this new species carries a diverse repertoire of secretion systems that is unique compared to all other *Xylella* spp. strains sequenced so far and might be involved in the interaction with the host plant and/or the unknown insect vectors.

In conclusion, this work greatly extends our knowledge of the genetic diversity, plant host range and distribution in natural environments of the genus *Xylella*. Despite being based solely on sequence data, our findings of new strains and even a new *Xylella* species in completely unsuspected areas (i.e. African savannah) and host plants such as wild rice opens new perspectives for future research and motivates surveillance programs in additional countries and biomes.

## DATA AVAILABILITY

The newly-assembled *Xylella* genomes described in this work are accessible at NCBI under BioProject PRJNA1241491.

## ACKNOWLEDGEMENTS

This work was funded by the Horizon Europe Project BeXyl (Beyond Xylella, Integrated Management Strategies for Mitigating Xylella fastidiosa Impact in Europe), Grant agreement no. 101060593. The authors also thank the Region Pays de la Loire as well as the bioinformatics platforms Genotoul and GenOuest for providing computational resources.

---

**Supplementary Table S1.** List of datasets in which the SRA Taxonomic Analysis Tool assigned more than 1,000 reads to *X. fastidiosa.* Run metadata, i.e. project type, target species, BioProject title and description are provided.

**Supplementary Table S2.** Sequences of *Xylella*-specific rRNA k-mers identified based on the SILVA SSU and LSU databases.

**Supplementary Table S3.** Blast identification of contigs assembled from *Xylella*-mapped reads from 21 datasets that did not contain *Xylella*.

**Supplementary Table S4.** List of contigs representing potential contaminants in published *X. fastidiosa* genomes.

**Supplementary Table S5.** List of 62 SRA datasets for which the presence of *Xylella* reads could be confirmed. Run metadata and k-mer counts for *Xylella* species/subspecies identification are provided.

**Supplementary Figure S1. KEGG pathway completeness in representative *Xylella* spp. genomes.** Heatmap showing the completeness of KEGG pathways/categories determined by KEGG Decoder. Only pathways detected in at least one strain are shown. *Xylella* species and *X. fastidiosa* subspecies are colour-coded as in Fig. 2. KEGG pathways/categories differentially represented in the four *X. rufipogonis* genomes are highlighted in blue.

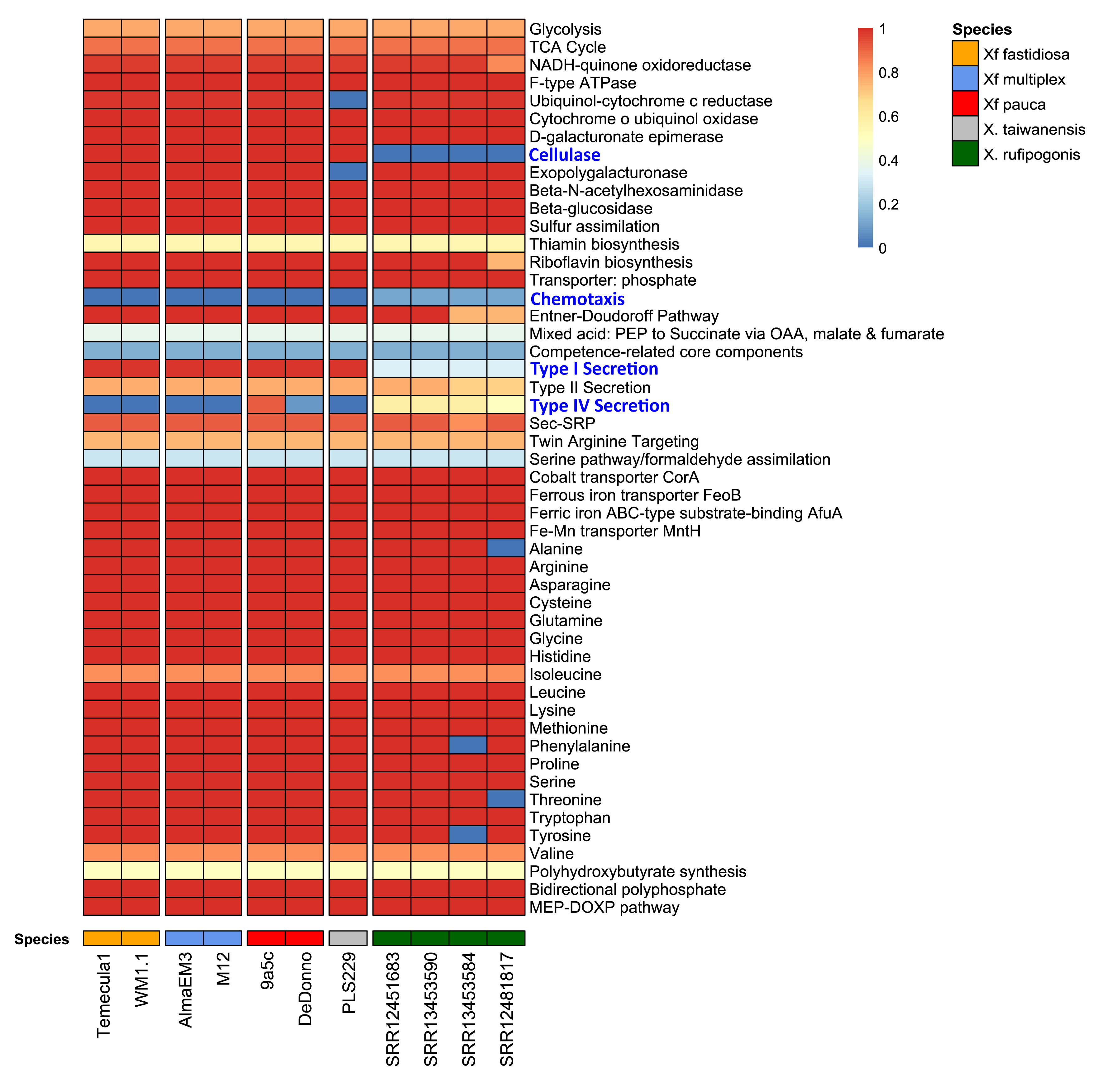

